# Visual stimulation drives retinotopic acetylcholine release in the mouse visual cortex

**DOI:** 10.1101/2024.02.04.578821

**Authors:** Scott G. Knudstrup, Catalina Martinez, Bradley C. Rauscher, Patrick R. Doran, Natalie Fomin-Thunemann, Kivilcim Kilic, John Jiang, Anna Devor, Martin Thunemann, Jeffrey P. Gavornik

## Abstract

Cholinergic signaling is involved with a variety of brain functions including learning and memory, attention, and behavioral state modulation. The spatiotemporal characteristics of neocortical acetylcholine (ACh) release in response to sensory inputs are poorly understood, but a lack of intra-region topographic organization of cholinergic projections from the basal forebrain has suggested diffuse release patterns and volume transmission. Here, we use mesoscopic imaging of fluorescent ACh sensors to show that visual stimulation results in ACh release patterns that conform to a retinotopic map of visual space in the mouse primary visual cortex, suggesting new modes of functional cholinergic signaling in cortical circuits.x

## Main

The neuromodulator acetylcholine (ACh) has widespread effects on cortical processes and can influence synaptic plasticity and neural dynamics (Colangelo et al., 2019; Hasselmo & Sarter, 2011; Picciotto et al., 2012). Although the importance of ACh in cortical function is widely appreciated, only recently has the development of fluorescent ACh indicators (Jing et al., 2018) made it possible to quantify cholinergic activity with spatiotemporal precision. A prominent view that behaviorally salient events evoke diffuse ACh release across the cortex has recently been challenged by imaging experiments that used these indicators to show heterogeneous patterns across cortical regions correlating with behavioral states and locomotion (Lohani et al., 2022). In the visual cortex, cholinergic signaling is involved in ocular dominance plasticity (Kasamatsu & Imamura, 2020) and required to encode spatiotemporal sequences (Gavornik & Bear, 2014) and intervals (Chubykin et al., 2013). While microdialysis has shown that visual stimulation can increase ACh levels in the visual cortex of anesthetized rats (Laplante et al., 2005), the extent to which this occurs during passive viewing in awake animals, and the spatiotemporal granularity of release, are unknown.

To address these questions, we performed a series of imaging experiments to measure ACh-dependent fluorescence across the dorsal cortical surface of the mouse cortex while awake animals passively viewed spatially localized visual stimuli (Fig. 1). Images were acquired using a four-color mesoscope capable of simultaneously measuring cholinergic, hemodynamic, and calcium activity described in Doran et al. (2023). A curved “crystal skull” window (Kim et al., 2016) allowed optical access to brain in mice (n=5) expressing the green ACh sensor GRAB_ACh3.0_ (Jing et al., 2020), three of which also co-expressed the red calcium indicator jRGECO1a (Dana et al., 2019).

**Fig 1.**
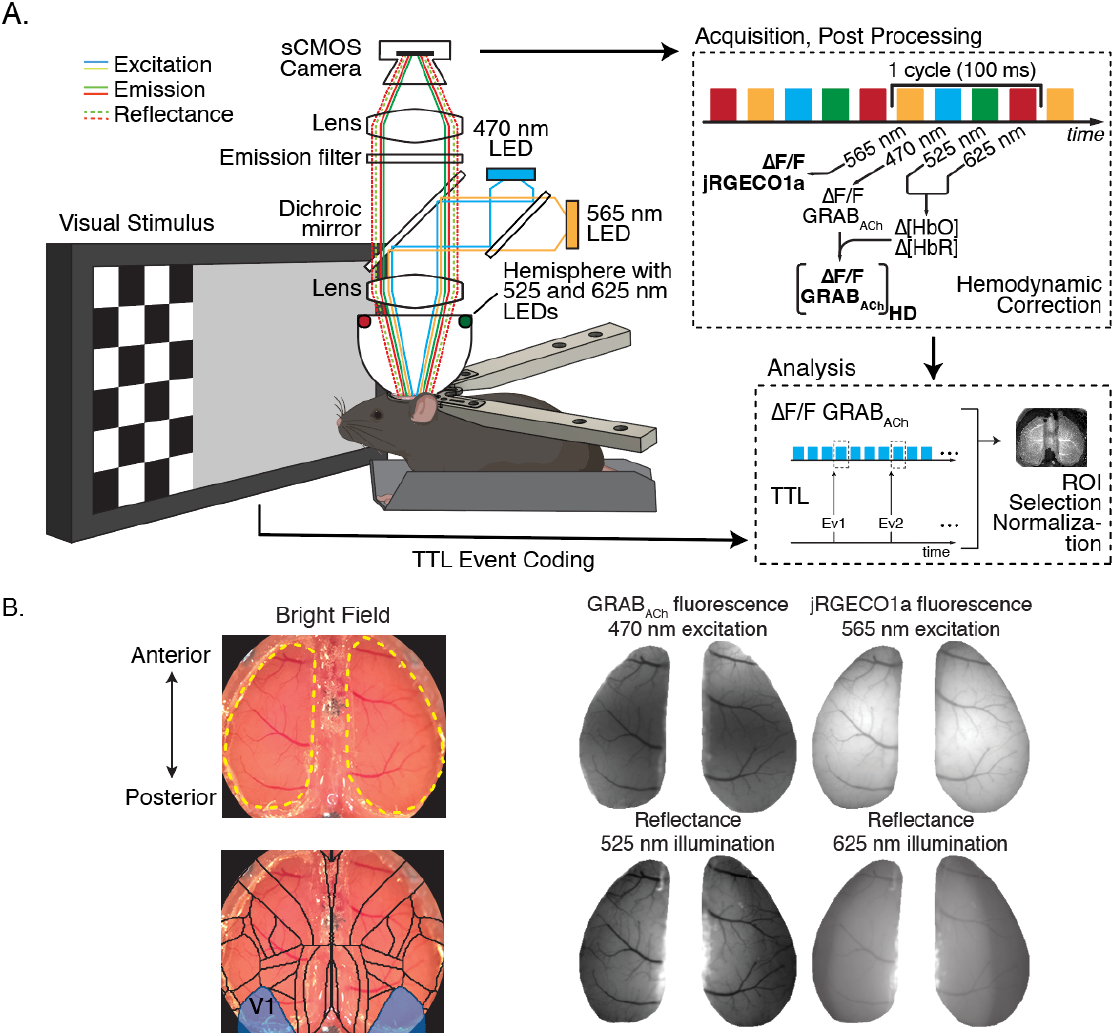
Mesoscopic imaging of ACh and Ca dynamics. **a**, Head-fixed, awake mice expressing either the GRAB_ACh_ sensor (n=2) or both GRAB_ACh_ and the jRGECO1a calcium indicators (n=3) passively viewed visual stimuli. A custom-built four-color mesoscope collected light from both fluorescent probes along with reflected light to estimate HbO and HbR concentration changes for hemodynamic correction of the GRAB_ACh_ fluorescence signals. Data were acquired at 10 Hz along with TTL-based event codes identifying visual stimuli within trials. After hemodynamic correction, ΔF/F timeseries extracted from regions of interest (ROIs) within visual cortical areas were analyzed relative to stimulus events. **b**, Example field-of-view showing the dorsal cortical surface visible to the mesoscope. Top: Left and right cortices are outlined (dashed yellow). Bottom: an overlay of the Allen Brain Institute Brain Atlas showing V1 (bottom left, shaded regions). **c**, Average intensity images of green (GRAB_ACh_) and red (jRGECO1a) fluorescence (top) as well as 525-nm and 625-nm reflectance (bottom).

Mouse eyes are wideset and most of the primary visual cortex (V1) represents contralateral monocular inputs from the peripheral visual field. To determine whether passive visual stimulus alone can drive measurable ACh release, and the extent to which this would be lateralized, we first presented mice expressing only the GRAB_ACh3.0_ sensor phase-reversing high-contrast checkerboard stimuli (Fig. 2a, left) located in peripheral regions of the left and right visual field. Stimuli were presented 50 times to each eye in 1 second blocks separated by 2 sec periods of static gray screen. Trial averaged fluorescence images (Fig. 2a, right) show that monocular visual stimulation alone is sufficient to drive a robust ACh increase that is restricted to the hemisphere contralateral to the viewing eye (see also supplemental video 1). The time course of ACh signaling in V1 differentiates between contralateral and ipsilateral hemispheres (Fig. 2b) and evoked ACh increase is significantly higher when viewing a patterned stimulus compared to gray screen viewing (Fig. 2c).

**Fig 2.**
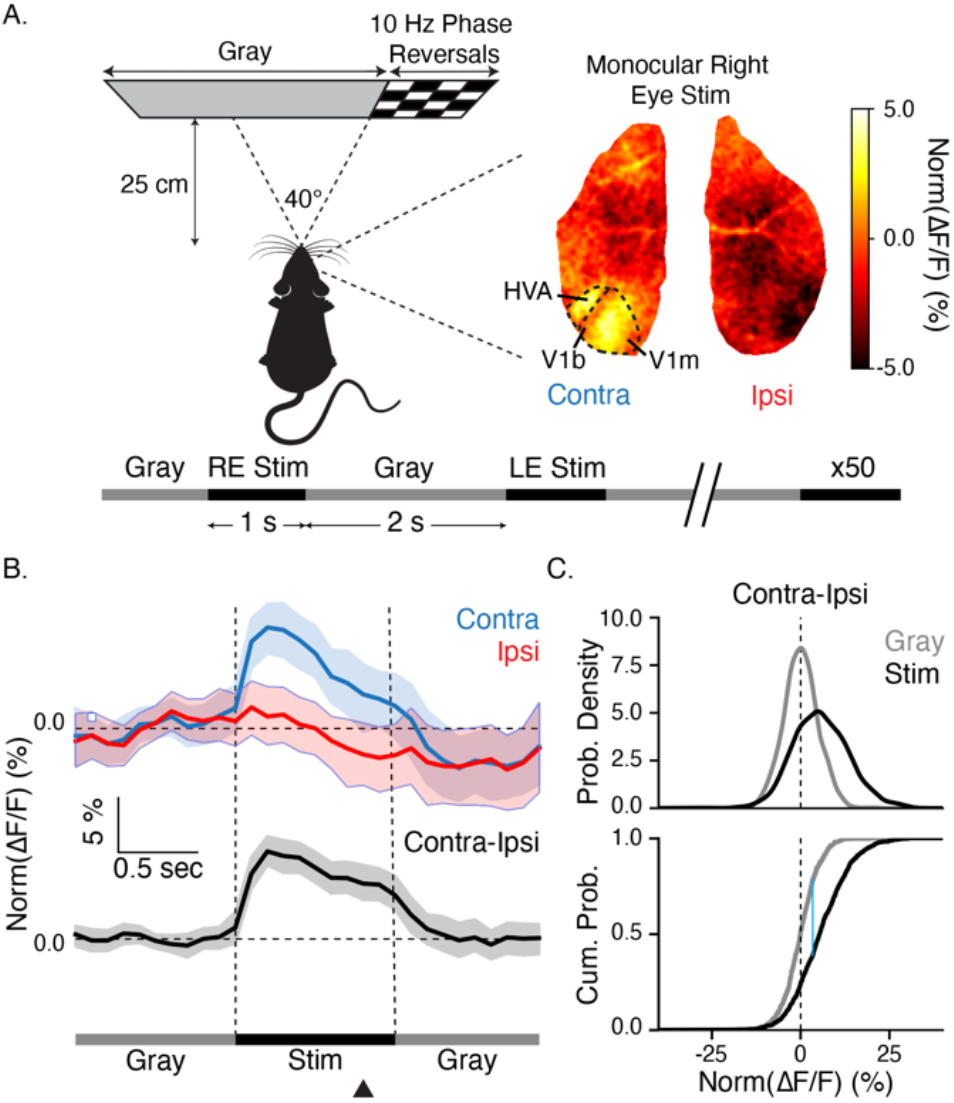
Monocular stimulation elicits lateralized ACh release localized to visual areas. **a**, Mice viewed a phase reversing checkerboard restricted to the left or right monocular regions of the visual field (left). Stimulus presentations lasted for 1 sec, alternated between the left and right visual field (50 trials each), and were separated by 2 secs of static gray screen. A trial-averaged GRAB_ACh_ fluorescence image (right) shows that monocular stimulation of the right eye elicits activity restricted to visual cortical areas in the left hemisphere (contralateral to the stimulus). Dashed lines demarcate approximate boundaries of V1 and higher visual areas (HVA). Binocular V1 (V1b) receives no visual stimulation and is relatively inactive compared to the monocular region (V1m). Fluorescence levels are shown as percent change from the baseline (ΔF/F) normalized by the total dynamic range calculated over the 300 sec acquisition period (see Methods) **b**, Trial-averaged GRAB_ACh_ fluorescence (top, shaded regions indicate 95% bootstrap confidence intervals around the mean) in V1m increases in the contralateral (blue), but not ipsilateral (red), cortex during stimulation. Accordingly, the difference between the two signals (black) is non-zero only during periods of stimulation demonstrating both stimulus-dependency and laterization of ACh release. The black triangle marks the location of the frame shown in panel A. **c**, Probability density (top) and cumulative distribution function (bottom, data corresponds to all frames in all trials for a given group, n = 100 trials x 10 frames = 1000) estimates show that the average difference between contra and ipsilateral V1m is centered at zero during gray periods but increases significantly with visual stimulation (KS-test, p = 5.4×10^-19^).

Cholinergic release was localized to the monocularly responsive region of V1 (V1m), matching stimulated peripheral areas of the visual field. Elevated fluorescence is also seen in neighboring higher visual areas (HVA, consisting of rostrolateral and anterolateral cortices, Fig. 2a) which are retinotopically organized but with azimuthal polarity reversed relative to V1 (Marshel et al., 2011). A band of relatively low activity separating V1m from the HVA demarcates binocular V1 (V1b) which in this experiment received inputs from the region of visual space occupied by a static gray screen. These results demonstrate that passive visual stimulation alone, absent any particular behavioral salience, can drive lateralized ACh release.

The surprising degree of localized fluorescence, closely matching the anatomical extent of cortex known to represent the stimulated regions of visual space, suggested that cholinergic release is retinotopically organized. To test this, we conducted a second experiment exposing mice expressing the GRAB_ACh_ sensor to a spatially precise stimulus consisting of a phase-reversing vertical checkered bar (Fig. 3a) that drifted from the monocular periphery into the binocular center of the visual field. ACh-dependent fluorescence was restricted to a strip of the right visual cortex (contralateral to viewing eye) that started in V1m and drifted laterally towards V1b as the bar moved across the visual field (Fig. 3b and supplemental videos 2-4). As the bar entered the binocular region, fluorescence was also seen ipsilateral to the viewing eye in V1b of the left hemisphere. Correlating the time of peak evoked fluorescence, which was about 1.5x higher than the average signal recorded simultaneously in ipsilateral V1m, with the stimulus azimuth produced a map showing a smooth retinotopic progression of ACh release across the cortical surface (Fig. 3c). Three of five mice in this experiment expressed both GRAB_ACh_ and the calcium sensor jRGECO1a, which allowed us to compare simultaneously acquired spatiotemporal dynamics of ACh and Ca signals. As expected from previous reports (Zhuang et al., 2017), the retinotopic stimulus produced a drifting strip of increased Ca activity (Supplemental Figure 3A). This Ca response closely matched that of ACh with comparable retinotopic maps calculated for both readouts (Fig. 3d, see also supplemental figure 3 and supplemental videos 2-5).

**Fig 3.**
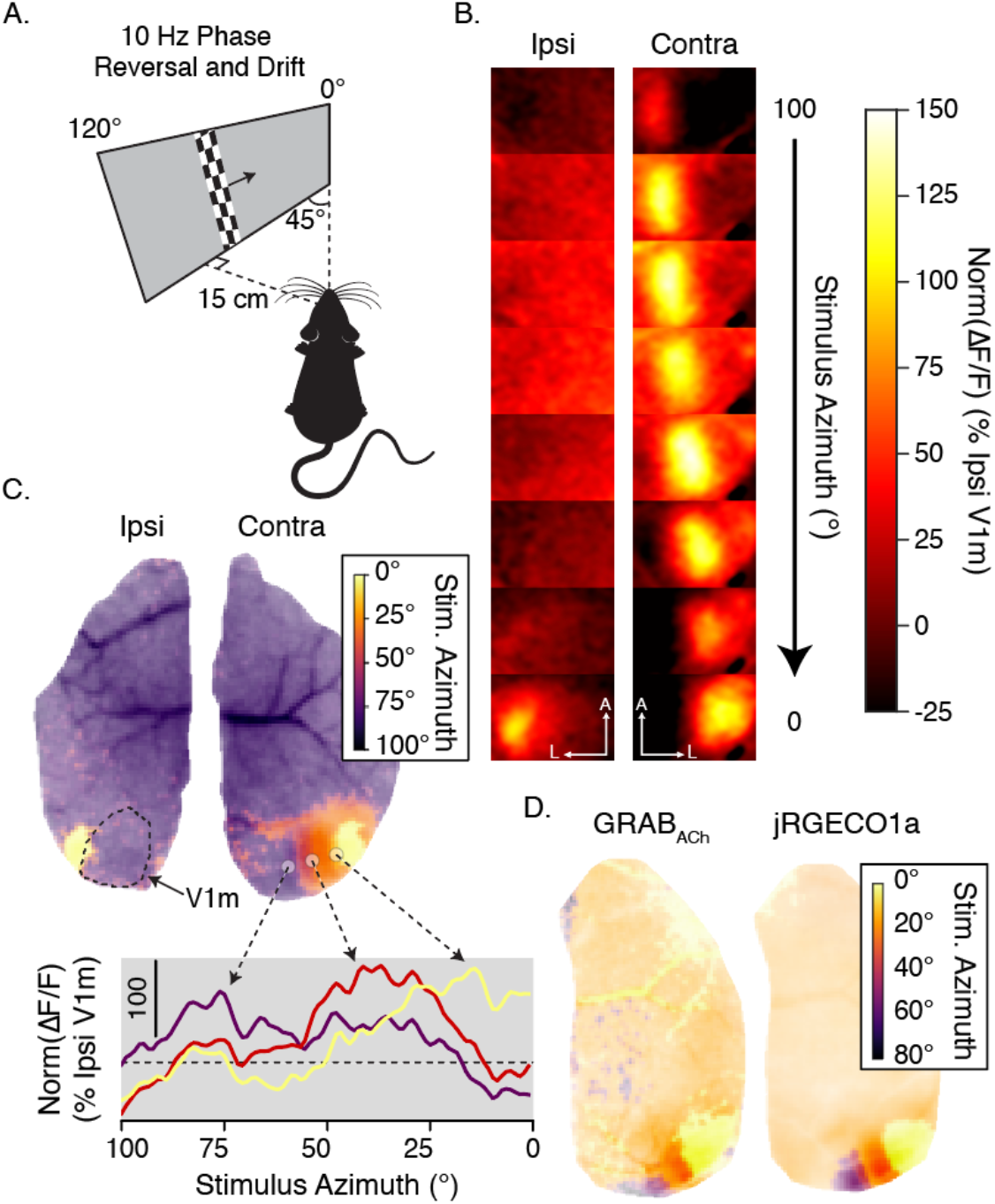
Acetylcholine release maps visual input space. **a**, A narrow, phase-reversing checkered bar swept across a screen positioned in the left visual field, traversing from the left periphery (∼120° off the centerline) to the center of the visual field (0°). **b**, Trial-averaged GRAB_ACh_ images (expressed as ΔF/F normalized by ipsilateral V1m, see Methods) for the ipsilateral (left) and contralateral (right, anterior/lateral directions marked in bottom frames) visual cortices as a function of stimulus azimuth showed a sweep of fluorescence across the cortex matching the progression of the visual stimulus. Increased fluorescence is observed in the ipsilateral hemisphere as the stimulus reaches the binocular region of the visual field (bottom-left image). **c**, A map (top) correlating the time of peak trial-averaged ACh signal with stimulus progression shows stimulus locked activity and retinotopic mapping in visual areas. Fluorescence (bottom) from three marked ROIs illustrates a pattern of spatially-specific ACh activity across the azimuthal axis of V1. **d**, ACh (left) and Ca (right) maps created in a mouse expressing both GRAB_ACh_ and jRGECO1a confirmed matching retinotopic maps for ACh and Ca.

## Discussion

Projections from the basal forebrain (BF) are the primary source of ACh in the visual cortex (Picciotto et al., 2012). Before the recent advent of fluorescent ACh sensors (Jing et al., 2018; Jing et al., 2020), it was difficult to quantify ACh release dynamics and there is scant evidence linking sensory input with precise patterns of cholinergic release. Though there is a long-standing debate about the extent to which cholinergic signaling occurs at individual synapses or through bulk volume transmission, a common theme in the literature is that ACh works to change the information processing state of broad neural ensembles (Colangelo et al., 2019; Muñoz & Rudy, 2014). This notion comports with our understanding of neuromodulators as working in a broad, distributed manner and is generally supported by studies tracing the anatomical distribution of cholinergic fibers. Indeed, a recent work using retrograde tracers found that while cholinergic BF projections into rat V1 and V2 originated from spatially segregated BF loci (Huppé-Gourgues et al., 2018), their innervation domains did not map onto retinotopically segregated locations within V1 (or HVAs) suggesting homogeneous ACh release within visual cortical areas.

Surprisingly, our functional imaging results showed that ACh release in primary and secondary visual areas was elicited by passive visual stimulation and produced patterns that formed a retinotopic map of visual space. Since we corrected ACh fluorescence for blood-oxygenation dependent light absorption, the observed cholinergic-dependent fluorescence dynamics are unlikely to be an artifact of the hemodynamic response. This mapping was observed in both mice expressing the GRAB_ACh_ sensor alone and in mice co-expressing the jRGECO1a Ca sensor, eliminating any concerns that the observed ACh signals could be an artifact resulting from crosstalk between the ACh and Ca channels.

There are at least two plausible explanations that might explain our findings. First, ACh could be released by cholinergic neurons in local cortical circuits. Specifically, a class of neurons expressing both choline acetyltransferase and vasoactive intestinal peptide (ChAT-VIP) target both interneurons and pyramidal cells across the cortical layers (Obermayer et al., 2019; Von Engelhardt et al., 2007) and might contribute to localized ACh release, though the functional role of ACh/ γ-Aminobutyric acid (GABA) co-release in these neurons remains unclear (Granger et al., 2016). Second, evoked cortical activity could induce ACh release from localized cholinergic axon collaterals independent of somatic activity in BF. Mechanistically, elevated potassium concentration microdomains resulting from visual activity (Singer & Dieter Lux, 1975) could directly result in axonal neurotransmitter release (Belhage et al., 1993; Contreras et al., 2021). In addition, there is a possibility that BF activity can selectively create topographic ACh release, though this would be at odds with the anatomical organization of efferent axons.

Localized ACh release conveys a variety of potential functional consequences. One implication is that we may need to supplement the notion of an externally directed “spotlight of attention” with local sources of “illumination” highlighting salient details based on feed-forward input patterns. The source of ACh in the cortex may vary between tonic and phasic activity patterns, with phasic ACh release driven by local cholinergic neurons automatically biasing cortical circuits to process feed-forward inputs (Hasselmo & Sarter, 2011) and wide-ranging tonic release driven by BF coordinating recurrent feed-back dominated oscillatory activity between cortical areas. Recent studies showing that ACh release in V1 increases neural activity in a layer-specific manner when paired with visuomotor stimulation (Yogesh & Keller, 2023), coordinates oscillatory rhythms in prefrontal areas (Howe et al., 2017), modulates prediction errors in the auditory cortex (Pérez-González et al., 2023), and can facilitate stimulus-induced learning (Yang et al., 2023) provide examples of how these dynamics can support behaviorally relevant function within local cortical circuits. Our experiments were restricted to the visual domain, and it will be important to determine whether similar release patterns occur in other sensory systems and brain regions. Regardless, these findings will require revisiting basic assumptions about how cholinergic signaling affects circuit dynamics, learning and memory and cortical plasticity.

## Methods

### Animals and surgery

All procedures were approved by the Institutional Animal Care and Use Committee (IACUC) of Boston University and followed institutional guidelines. Mice were housed in a climate-controlled environment on a standard 12-hour light-dark cycle and were provided with food and water *ad libitum*. Experiments were performed during the mouse’s light cycle. Five mice aged 2-5 months were used in this study. All five animals expressed GRAB_ACh_, and three of five mice carried the Thy1-jRGECO1a(GP8.20)Dkim transgene and expressed the calcium indicator jRGECO1a (Dana et al., 2016, 2019). To induce cortex-wide GRAB_ACh_ expression, 1.5 μL of AAV9-hSyn-GRABACh3.0 (WZ Biosciences, 3.06×10^13^ GC/mL) was injected into each transverse sinus (to total of 3 μL per animal) of 1-day old neonates from a [Thy1-jRGECO1a/wt] x [wt/wt] breeding following procedures described in Lohani et al. (2022). Headpost implantation and craniotomy were performed during a single surgery in adult (8–16-week-old) animals following procedures described in Kılıç et al. (2021). A custom-designed, titanium headbar was attached to the cranium and a modified crystal skull glass window was used to replace the dorsal cranium over both hemispheres; the original curved glass (width: 12 mm, labmaker.org) was cut in half to obtain separate glass pieces for each hemisphere. Following the implantation procedure, a silicone plug was placed on top of the glass window and a 3D-printed cap was fixed to the headpost to prevent heat loss. Mice could recover for at least one week after surgery before behavioral training commenced. Animals were head-fixed for increasing periods while receiving rewards of sweetened condensed milk. Data acquisition started when animals tolerated one hour of head fixation.

### Visual stimulation

Visual stimuli were generated and displayed using the PsychToolbox extension for MATLAB, and custom software was used to stimulus control timing and hardware signals. Stimuli were displayed on a gamma corrected 27-inch LED monitor (ASUS VG279Q, 1920 x 1080 pixels, 120 Hz refresh rate) and matched for uniform average luminance.

Monocular stimulation consisted of a phase reversing checkerboard pattern (black/white phase reversals every 100 ms, or 10 Hz) presented outside of the binocular center of the visual field. The checkerboard stimulus occupied a region of the visual field ranging from 20°-63°. The central/binocular region was a uniform gray throughout as was the non-stimulus half of the screen (e.g. when stimulus was being shown in the left visual field, the center and right side of the screen were both static gray). Stimulation periods lasted 1 sec each and were separated by 2 sec of gray screen. Stimulation alternated between the left-and right-side (n=50 each). The screen was placed 25 cm directly in front of the mouse.

For the retinotopic mapping experiment, the screen was positioned 15 cm measured tangentially from the animal’s left eye and angled 45° relative to the mouse’s midline. The stimulus consisted of a narrow vertical bar (3° wide) that swept from 120° (left periphery) to 0° (center) of the visual field on a solid gray background. The bar resembled a narrow section of a checkerboard with squares 5° wide that phase reversed at 10 Hz. To keep the width (in degrees) and spatial frequency of the stimulus constant as it swept across the screen, we applied spherical correction to render the stimulus in spherical coordinates (Marshel et al., 2011). Sweeps lasted 11 s (n=100) and were presented in immediate succession (i.e., no gray screen between trials).

### Mesoscopic imaging

Fluorescent images were acquired with a custom widefield imaging system using a single sCMOS camera (Sona 4.2B-6, Andor) for sequential recording of GRAB_ACh3.0_ and jRGECO1a fluorescence as well as hemoglobin absorption at 525-nm and 625-nm (Doran et al. 2023). The field of view covered a 10×10 mm^2^ area containing the entire dorsal cortical surface, and each channel was captured at 10 Hz. The spatial resolution was 20 μm at 4×4 binning. For fluorescence excitation, epi-illumination was delivered through the objective lens (ZDM-1-MVX063, Olympus). For GRAB_ACh3.0_ excitation, we used a 470-nm LED (SOLIS-470C, ThorLabs) filtered through a 466/40 nm bandpass filter (Semrock). For jRGECO1a, we used a 565-nm LED (SOLIS-565C, ThorLabs) filtered through a 560/14-nm bandpass filter (Semrock) and an additional 450-nm longpass filter (Chroma). Emitted light passed through a 488/561-nm dual-band dichroic mirror (Semrock) and a multi-band 523/610-nm filter (Semrock) which allowed GRAB_ACh3.0_ and jRGECO1a fluorescence (500-540 nm and 580-640 nm, respectively) to reach the camera while blocking excitation light at 440-480 nm and 550-570 nm. LEDs for hemoglobin absorption were placed in a custom-made reflectance illumination ring between the objective lens and sample surface. To prevent illumination light from reaching the mouse’s eye and causing unwanted visual stimulation, an aluminum hemisphere with a ∼1.5-cm opening at the bottom was placed between objective lens, reflectance illumination ring, and the surface of the cranial window preparation. See Doran et al. (2023) for a detailed description of the imaging system.

### Data preprocessing

A baseline image was calculated for each channel by taking the temporal average of the entire time series. For fluorescence channels, each image was divided by the baseline image to calculate ΔF/F. The 525-nm and 625-nm reflectance channels were used to estimate oxy- and deoxyhemoglobin concentration changes, which were then used to correct the green fluorescence signal for intensity changes due to absorption of fluorescence excitation and emission light by hemoglobin, as described in Ma et al. (2016).

### ROI extraction and time series creation

we used trial-averaged videos and the Allen mouse brain atlas (Wang et al., 2020) to locate left and right V1 and higher visual areas. To obtain trial-averaged videos, we collected all frames following stimulus onset (10 frames for monocular stimulation, 110 frames for retinotopic stimulation) and averaged across trials for each timepoint. Thus, frame *i* in a trial-averaged video represents the average of all trials’ *i*^th^ frame. Timeseries for individual ROIs as specified in the results section were computed by averaging over all pixels within the ROI for each frame with normalization described below.

### Monocular stimulation analysis

After extracting timeseries for left and right V1, data were high-pass filtered (scipy.signal.butter, cutoff=0.1 Hz, order=2) to remove slow oscillations (see Figure S1) and we measured the full range of the filtered data. We next extracted a 3 s period around each trial (1 s pre-stimulus/baseline gray screen, 1 s stimulus, 1 s post-stimulus gray screen), computed the mean activity during the pre-stimulus baseline period, subtracted this value from all data points in the 3 second window, and renormalized the time series as a percent of total range, e.g. 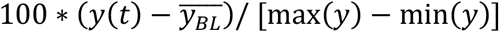 where *y*(*t*) is the high-pass filtered timeseries after hemodynamic correction, max & min(*y*) define the total dynamic range within the ROI over the entire experiment, and 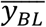 is the average fluorescence during the pre-stimulus baseline period. Norm(ΔF/F) (%) values represent trial averaged rescaled fluorescence as a percent of the ROI’s range over the recording session. Trials were combined into contralateral and ipsilateral groups (e.g., right V1 during left-eye stimulation was combined with left V1 during right-eye stimulation to produce the contralateral group) and bootstrap trial mean estimates with 95% confidence intervals were calculated for both groups. The difference between contralateral and ipsilateral responses was calculated by subtracting the ipsilateral response from the contralateral response on a trial-by-trial basis before calculating the trial-averaged mean and bootstrapped confidence intervals.

To assess the statistical significance of enhanced contralateral responses relative to ipsilateral response, we compared contra-ipsi values for all timepoints in pre-stimulus baseline and stimulation periods (100 trials x 10 frames = 1000 samples per group). We used the python functions scipy.stats.gaussian_kde and scipy.stats.ecdf to estimate empirical probability density and cumulative density functions (PDF and CDF) and scipy.stats.ks_2samp to perform a two sample KS-test on this data.

### Retinotopic mapping

To produce a retinotopic map, we first produced a trial-averaged video of ACh and Ca fluorescence images and applied light gaussian smoothing (kernel sizes: time = 50 ms, x and y = 1 pixel) to reduce noise. For each pixel, we found the location in time of its peak average fluorescence and converted this value into degrees azimuth based on the location of the sweeping bar in the visual field at that time. These values were mapped into color space. To emphasize the strength of pixel modulation, we used the magnitude peak fluorescence to set color opacity in resulting maps. Our imaging window does not include the entirety of V1 and so we are unable to image fluorescence including the full range of mouse vision (∼180° compared to ∼100° in the mouse expressing only GRAB_ACh_ and ∼80° in the mice with co-expressing jRGECO1a).

To produce activity traces at different spatial location across the cortical surface we selected three 5×5 pixel patches across V1’s retinotopic axis and produced a time series for each by averaging over all pixels within each patch for each frame in the trial-averaged video. To compare the fluorescence in a localized region receiving visual inputs with baseline activity in a region that was not being stimulated we rescaled these traces based on the activity in ipsilateral monocular V1 (V1m). Specifically, we drew an ROI around ipsilateral V1m, extracted its time series as above and measured the dynamic range of total fluorescence. The rescaled time series, labeled as Norm(ΔF/F) (% ipsi V1m), reflects stimulated fluorescence as a percent of unstimulated fluorescence, calculated as 100 * *y*(*t*)/[max (*V*1*m*_*ipsi*_) *−* min (*V*1*m*_*ipsi*_)], where *y*(*t*) is the average hemodynamic corrected time series from a small patch and & min (*V*1*m*_*ipsi*_) define the range of values from ipsilateral V1m over the trial averaged time series. The same rescaling method was also applied to all pixels in Supplementary retinotopy videos 2-5.

## Acknowledgements

This work was supported by NEI R001EY030200, NIH/Brain Initiative U19NS123717, NIDA R01DA050159, and NIH/Brain Initiative R01NS122742.

---

**Fig S1.**
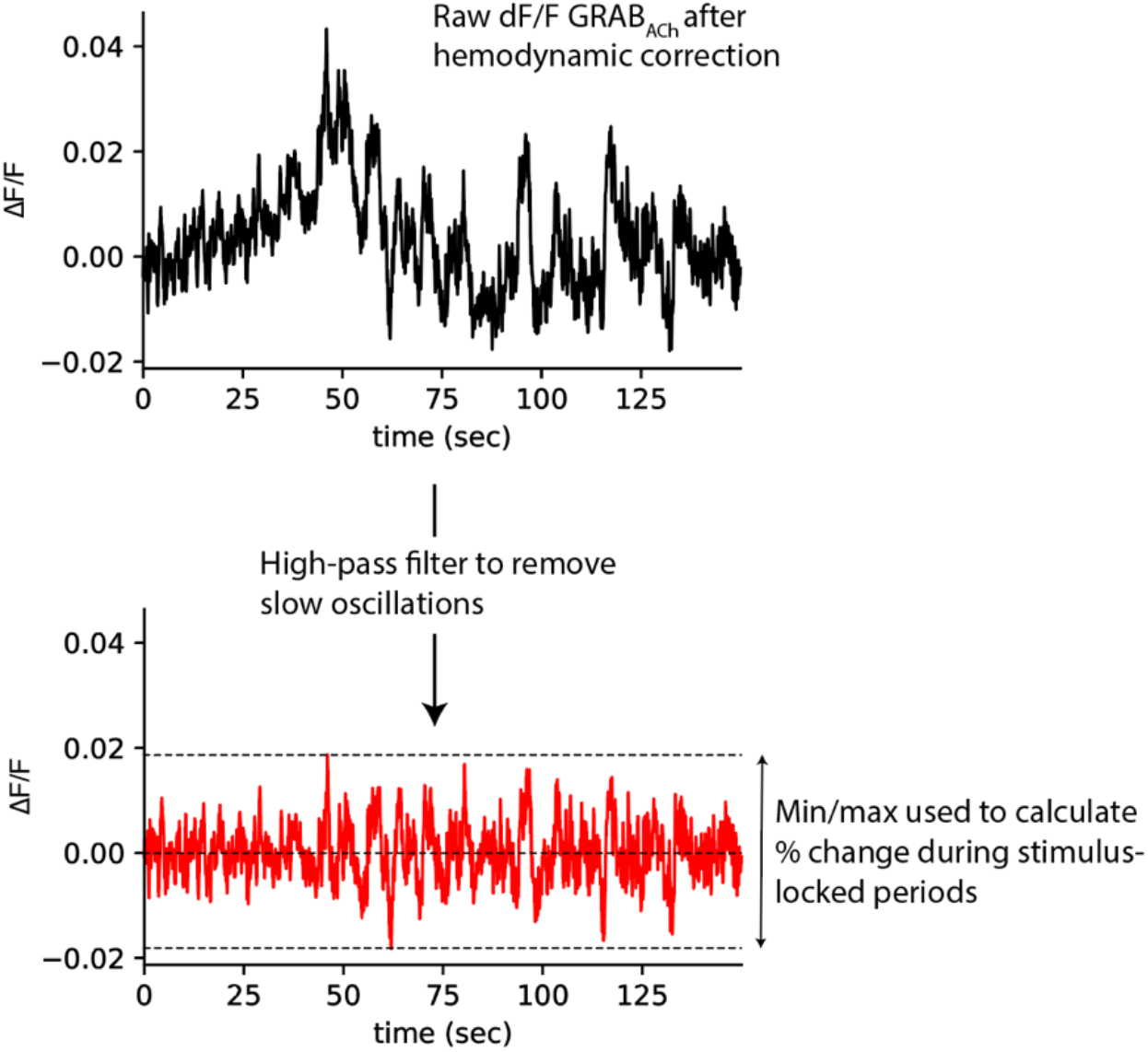
High-pass filtering. Example ΔF/F data from from left V1 after hemodynamic correction (top, 150 sec shown from ∼6 minutes of imaging during the monocular stimulation experiment). Slow fluctuations increase the total dynamic range of the data. To define a functional range that could be used to compare stimulus evoked fluctuations at different points in time, we used a high-pass filter (Butterworth, cutoff = 0.1 Hz, order=2) to remove slow oscillations. The functional range from filtered data (bottom) was used to calculate percent changes of stimulus evoked activity relative to gray-screen baseline measurements on a trial-by-trial basis as detailed in the methods section.

**Fig S2.**
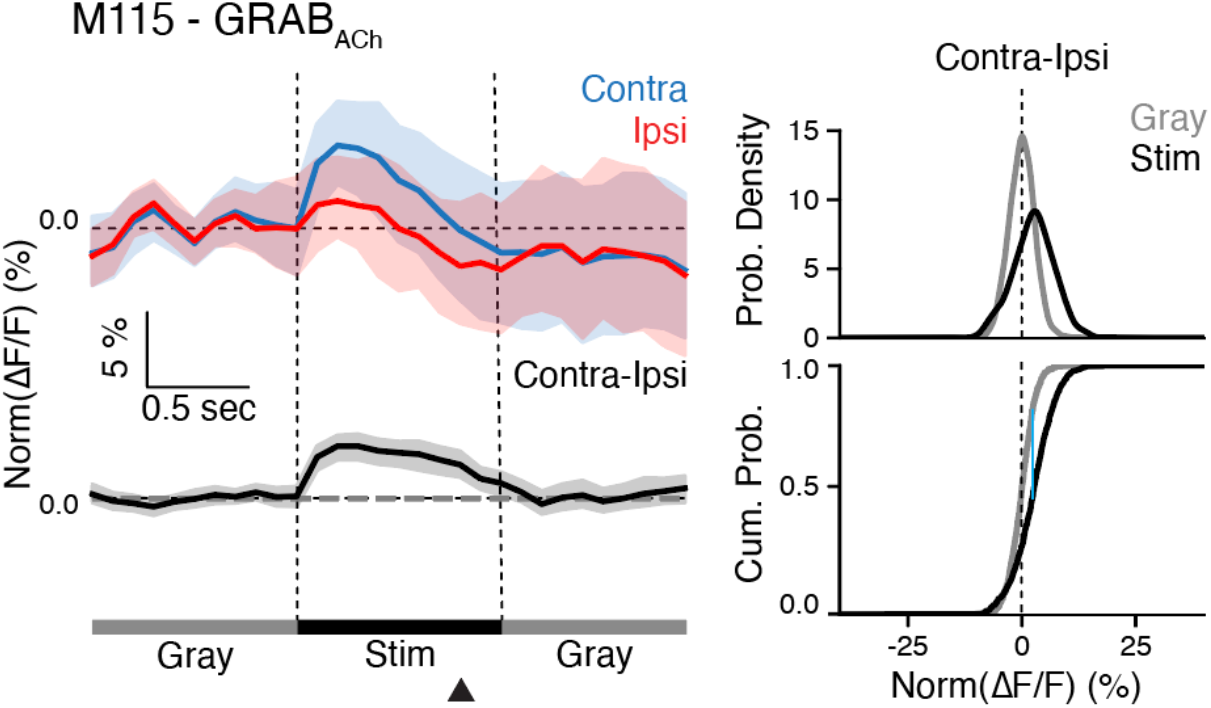
Monocular stimulation for mouse M115. Analysis of fluorescence from a second mouse expressing only the GRAB_ACh_ sensor (i.e., no expression of jRGECO1a) demonstrates hemisphere-specific cholinergic release with monocular stimulation. Analysis as in Fig. 2b-c. Fluorescence in this mouse was weaker than in the other mouse (the source for Fig 2), likely due to differences in cranial window clarity or GRAB_ACh_ expression level, but the difference between contralateral and ipsilateral responses to stimulation show the same statistically significant effect of monocular stimulation (KS-test; n=1000, p = 2.5×10^-13^).

**Fig S3.**
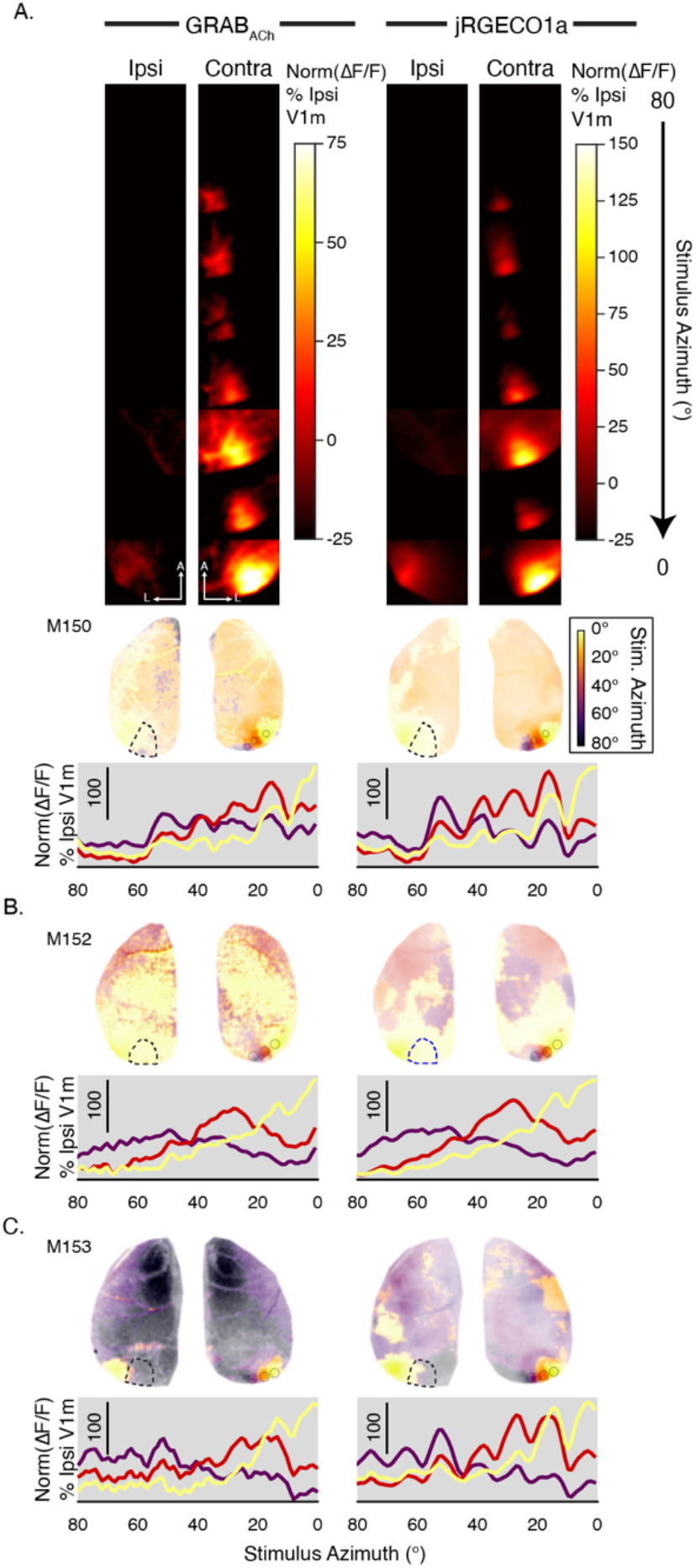
Retinotopic maps from mice co-expressing GRAB_ACh_ and jRGECO1. **a**, Fluorescence time-courses (top) and retinotopic maps with fluorescence time series (bottom) from a mouse co-expressing GRAB_ACh_ and jRGECO_1a_ (M150, some data also shown in Figure 3D). **b-c**, Retinotopic maps calculated for two additional co-expressing mice (M152 and M153). In all panels the left column shows data from the GRAB_ACh_ channel and the right from jRGECO_1a_. The stimulus azimuth color code is the same for all three mice. All data analysis conducted and presented as in Figure 3.

**Supplemental Video 1 | Monocular stimulation (GRAB_ACh_ only)** Video accompanying Fig. 2 showing trial-averaged GRAB_ACh_ fluorescence during monocular left-eye (left panel) and right-eye (right panel) stimulation, including 1s pre- and post-stimulus periods (3 sec, or 30 frames total). The video was scaled (see Methods) such that Norm(ΔF/F) (%) values represents fluorescence as a percent of dynamic range over the entire recording session.

**Supplemental Video 2** | **Retinotopic stimulation (GRAB**_**ACh**_ **only)**. Video accompanying Fig. 3b/c showing trial-averaged GRAB_ACh_ fluorescence during drifting bar stimulation. Stimulus position is indicated at the top left. Fluorescence is displayed as a percentage of the dynamic range in ipsilateral monocular V1 (see Methods).

**Supplemental Videos 3-5** | **Retinotopic stimulation (jRGECO1a and GRAB**_**ACh**_). Videos accompanying Fig. 3d and S3 showing simultaneous trial-averaged jRGECO1a and GRAB_ACh_ fluorescence from three separate mice during drifting bar stimulation. Stimulus position is indicated at the top left. Fluorescence is displayed as a percentage of the dynamic range in ipsilateral monocular V1 (see Methods). Videos S3-S5 correspond to mice M150, M152, and M153 respectively.

